# The Dynorphin/Kappa Opioid Receptor mediates adverse immunological and behavioral outcomes induced by repetitive blast trauma

**DOI:** 10.1101/2022.08.15.504055

**Authors:** Suhjung Janet Lee, Aric F. Logsdon, Mayumi Yagi, Britahny M. Baskin, Elaine. R. Peskind, Murray M. Raskind, David G. Cook, Abigail. G. Schindler

**Author notes:** Corresponding Author: Abigail G. Schindler, PhD, VA Puget Sound Health Care System, S182, 1660 South Columbian Way, Seattle, WA 98108.

## Abstract

**Background:** Adverse pathophysiological and behavioral outcomes related to mild traumatic brain injury (mTBI), posttraumatic stress disorder (PTSD), and chronic pain are common following blast exposure and contribute to decreased quality of life, but underlying mechanisms and prophylactic/treatment options remain limited. The dynorphin/kappa opioid receptor (KOR) system helps regulate behavioral and inflammatory responses to stress and injury; however, it has yet to be investigated as a potential mechanism in either humans or animals exposed to blast. We hypothesized that blast-induced KOR activation mediates adverse outcomes related to inflammation and affective behavioral response.

**Methods:** C57Bl/6 adult male mice were singly or repeatedly exposed to either sham (anesthesia only) or blast delivered by a pneumatic shock tube. The selective KOR antagonist norBNI or vehicle (saline) was administered 72 hours prior to repetitive blast or sham exposure. Serum and brain were collected 10 minutes or 4 hours post-exposure for dynorphin A-like immunoreactivity and cytokine measurements, respectively. At one-month post-exposure, mice were tested in a series of behavioral assays related to adverse outcomes reported by humans with blast trauma.

**Results:** Repetitive but not single blast exposure resulted in increased brain dynorphin A-like immunoreactivity. norBNI pretreatment blocked or significantly reduced blast-induced increase in serum and brain cytokines, including IL-6, at 4 hours post exposure and aversive/anxiety-like behavioral dysfunction at one month post exposure.

**Conclusions:** Our findings demonstrate a previously unreported role for the dynorphin/KOR system as a mediator of biochemical and behavioral dysfunction following repetitive blast exposure and highlight this system as a potential prophylactic/therapeutic treatment target.

## INTRODUCTION

Mild traumatic brain injury (mTBI) is a serious public health issue for Veterans and military Servicemembers (SM). Moreover, proliferating use of high explosives against civilian populations in regions of conflict pose growing risks of blast-related mTBI globally. Approximately 75-90% of all traumatic brain injuries in the United States are mild (CDC 2003; WHO 2006; DOD 2021). Often referred to as the “signature injury” in Veterans and SMs from the Iraq/Afghanistan wars (OIF/OEF/OND) (O’neil et al., 2013), mTBI is reported at higher incidence in SMs compared to the civilian population (Reid & Velez 2015). While approximately 10-20% of individuals from OEF/OIF/OND have experienced at least one TBI (Tanielian and Jaycox 2008; Lindquist 2017), 75-85% of all TBIs reported are from repetitive blast exposure (Hoge et al., 2008; Terrio et al., 2009). Likewise, combat training, terrorist bombings, and industrial accidents are also commonly associated with blast trauma (Rosenfeld et al., 2013). While health issues can resolve within several weeks, significant persisting symptoms (cognitive, affective, and pain interference) can decrease quality of life and daily functioning (McMahon et al., 2014; Agtarap et al., 2021). Critically, preventive measures and treatment options for these chronic and debilitating symptoms remain limited.

Mechanisms underlying adverse outcomes following repetitive blast exposure are not understood and are an area of active investigation. Roughly 40% of Veterans from OIF/OEF/OND with a history of blast exposure met criteria for polytrauma (e.g., the Polytrauma Clinical Triad), consisting of post concussive symptoms from mTBI, posttraumatic stress disorder (PTSD), and chronic pain (Lew et al., 2009). Results from animal studies also demonstrate comorbid outcomes related to polytrauma following blast exposure. We and others have reported on similar behavioral phenotypes in rodents following blast exposure as is present in SMs and Veterans with a history of blast trauma, including disinhibition and risk taking (Schindler et al., 2017), executive dysfunction (Baskin et al., 2021), substance use disorder risk (Schindler et al., 2021b), and aversion/increased stress-reactivity (Schindler et al., 2021a; Perez-Garcia et al., 2016; Elder et al., 2012).

One potential molecular mechanism capable of contributing to both mTBI, PTSD, and chronic pain-related outcomes is activation of the dynorphin/kappa opioid receptor (KOR) system. Indeed, this receptor system has been implicated in adverse outcomes following mild to severe head impact trauma (Hauser et al., 2005; Pietrzak et al. 2014; Best et al., 2022), but has not been investigated in a blast trauma setting (mild or severe). KORs are expressed widely throughout the brain and dynorphin/KOR activation is responsible for the dysphoric/aversive component of stress and pain (Bruchas et al., 2010; Land et al., 2008, Cahill et al., 2014). Likewise, mild blast trauma is associated with hypothalamic-pituitary-adrenal (HPA) axis dysfunction (Wilkinson et al., 2012) and KORs are expressed within the HPA axis where they help regulate stress response and neuroendocrine function (Van’t Veer & Carlezon 2013). Cortisol levels are acutely increased in human males post blast exposure (Cernak at al., 1999) and we demonstrated a similar blast-dose effect on increased corticosterone release in male mice (Schindler et al., 2021a). In that study, we also demonstrated that repetitive but not single blast exposure results in aversion to blast-paired cues, highlighting the potential for dynorphin/KOR to play a crucial role in adverse blast outcomes.

Dysregulation of the immune system is also a potential mechanism underlying adverse blast outcomes and may interact with the dynorphin/KOR system. Both TBI and PTSD alone can result in increased production of inflammatory cytokines such as IL-6 and TNF-α (Ziebell et al., 2010; Speer et al., 2018; Deslauriers et al., 2017). Likewise, in SMs and Veterans with comorbid PTSD and TBI, IL-6 increase is associated with symptom severity (Gill et al., 2014; Rodney et al., 2020) and is considered a therapeutic target of interest (Monsour et al., 2022). Additionally, HPA axis dysfunction in mTBI may be linked to altered pro- and anti-inflammatory responses (Tapp et al., 2019). KORs are expressed on immune cells (Machelska & Celik 2020), can influence IL-6 and TNF-α levels (Alicea et al., 1995), and help mediate aspects of acute and chronic neuropathic pain (Cahill et al., 2014; Liu et al., 2019; Massaly et al., 2019). While previous results have demonstrated acute dynorphin increase following mild to severe impact-induced brain injury in cats and rodents (McIntosh et al., 1987; Redell et al., 2002; Hussain et al., 2012), the dynorphin/KOR system has not been studied in blast-exposed individuals nor in animal blast models.

These considerations lead us to hypothesize that repetitive mild blast exposure results in dynorphin level increase and subsequent activation of the KOR system to mediate adverse outcomes following blast exposure. To test this, we utilized our well-established mouse model of blast that accurately recapitulates open-field blast forces (Meabon et al., 2016; Logsdon et al., 2018; Logsdon et al., 2020; Schindler et al., 2017; Schindler et al., 2021a; Schindler et al., 2021b; Baskin et al., 2021) in combination with established dynorphin A-like immunoreactivity and cytokine assays and behavioral tests related to adverse blast outcomes. Building on our finding of increased serum corticosterone acutely following blast exposure (Schindler et al., 2021a), here we show that repetitive but not single blast exposure resulted in an acute increase in brain dynorphin A-like immunoreactivity of male mice. We then tested the ability of a KOR antagonist to block and/or reduce adverse blast outcomes by pre-treating male mice with norbinaltophimine (norBNI), a selective long-acting (receptor inactivating) KOR-antagonist lasting up to 21 days in rodents (Bruchas & Chavkin, 2010; Bruchas, Land, & Chavkin, 2010; Bruchas et al., 2007; Chavkin, Cohen, & Land, 2019; Schattauer et al., 2017). Pretreatment with norBNI prior to repetitive blast exposure blocked some, but not all, acute cytokine changes in both blood serum and brain. Importantly, we further demonstrate that KOR antagonism completely blocked aversive- and anxiety-like behavioral outcomes and blast-induced light aversion but not motor incoordination. Taken together, these data suggest that the dynorphin/KOR system mediates adverse blast-induced inflammatory response and behavioral affective regulation.

## MATERIALS AND METHODS

### Animals

Male wild-type C57Bl/6 mice (Jackson Laboratory) 9 weeks of age upon arrival were given one week to acclimate, followed by an additional week of handling habituation prior to sham or blast exposure. Animals are randomly assigned by cage (3 animals per cage) to vehicle, drug, sham, or blast groups. Animals were group-housed with access to food and water ad-libitum on a 12:12 light:dark cycle (lights on at 6 am), with the exception of during training for the photophobia assay where their food was restricted to maintain 83%-90% of their ad libitum body weight. All animal experiments were conducted in accordance with Association for Assessment and Accreditation of Laboratory Animal Care guidelines and were approved by the VA Puget Sound Institutional Animal Care and Use Committee.

### KOR antagonist pretreatment

Three days prior to sham/blast exposure, animals were injected intraperitoneally with either 10 mg/kg nor-binaltorphimine dihydrochloride (EMD Millipore, Burlington, MA) dissolved in sterile saline or vehicle (saline; 10 mL/kg).

### Mouse model of blast trauma

The shock tube (Baker Engineering and Risk Consultants, San Antonio, TX) was designed to generate blast overpressures (BOP) that mimic open field high explosive detonations experienced by military SMs in the Iraq and Afghanistan wars. Design and modeling characteristics have been described in detail elsewhere (Schindler et al., 2017, Huber et al., 2013). Briefly, mice were anesthetized with isoflurane (induced at 5% and maintained at 2%), secured to gurney, and then placed into shock tubes where the ventral body surface was facing the oncoming shock wave (Koliatsos et al., 2011). Sham control animals received anesthesia only (same room and induction chamber as for blast exposure) for a duration matched to blast animals. Anesthesia duration ranged from 3-5 minutes. Animals were exposed to sham/blast exposure either once (1x) or over the course of three days, one per day (3x). Following exposure, mice were immediately removed from the shock tube, anesthesia discontinued, and the mouse placed on its back to determine loss of righting reflex (LORR) time (amount of time required for the mouse to flip over and place hind legs 2x). Average BOP peak intensity (psi) was in keeping with mild blast TBI (19.59 psi +/- 1.41) (Koliatsos et al., 2011, Cernack et al., 2011). Under these experimental conditions, the overall survival rate is 97%, with blast-exposed mice comparable to sham-exposed mice upon inspection 2-4 h following exposure (e.g., responsive to stimuli, normal posture, and breathing). Animals were weighed on each day of blast/exposure, 24-, 48-hours, and 1-month post-exposure. Sham 1x and 3x animals were pooled together as there were no statistically significant differences for the dynorphin immunoassay.

### Enzyme-Linked Immunosorbent Assay, Immunoassays

#### Cytokine

Animals (n’s: sham vehicle: 8, sham norBNI: 12, blast vehicle: 9, blast norBNI: 12) were sacrificed 4 hours following final sham/blast exposure and trunk blood and brain collected for downstream biochemical analysis (Figure 1A). Serum was collected in a serum separator gel tube and was allowed to clot at room temperature for 30-45 minutes before centrifugation (3000 x g) for 10 min. Brains were placed in a 2.0 mL BeadBug pre-filled microcentrifuge tube on ice containing 1.0 mm acid washed silica beads (Sigma, Burlington, MA) and 1 mL of total lysis buffer solution. Lysis buffer consisted of 0.02% Triton-X, 10mM HEPES, 1.5mM MgCl2, and 10mM KCl, with fresh protease/phosphatase inhibitor cocktails (Sigma, Burlington, MA) (Erickson & Banks, 2011). Brains in ice cold lysis buffer were then homogenized twice with a BeadBug (Sigma, Burlington, MA) at 4000 rpm for 45 seconds each The samples were then centrifuged at 4°C at 18,000 x g for 10 minutes and the supernatant was aliquoted and stored at −80°C. Pro- and anti-inflammatory cytokine level were then analyzed using the IDEXX Bioanalytics Mouse Multiplex Cytokine Panel (23-plex) and service (samples shipped to IDEXX on dry ice overnight). To normalize variable protein amounts, brain lysates were assayed for total protein concentration using Pierce BCA protein assay (Thermo Fisher Scientific, Rockford, Illinois) as per manufacturer protocol (50-fold dilution). Measurements were completed by reading absorbance at 562 nm.

**Figure 1.**
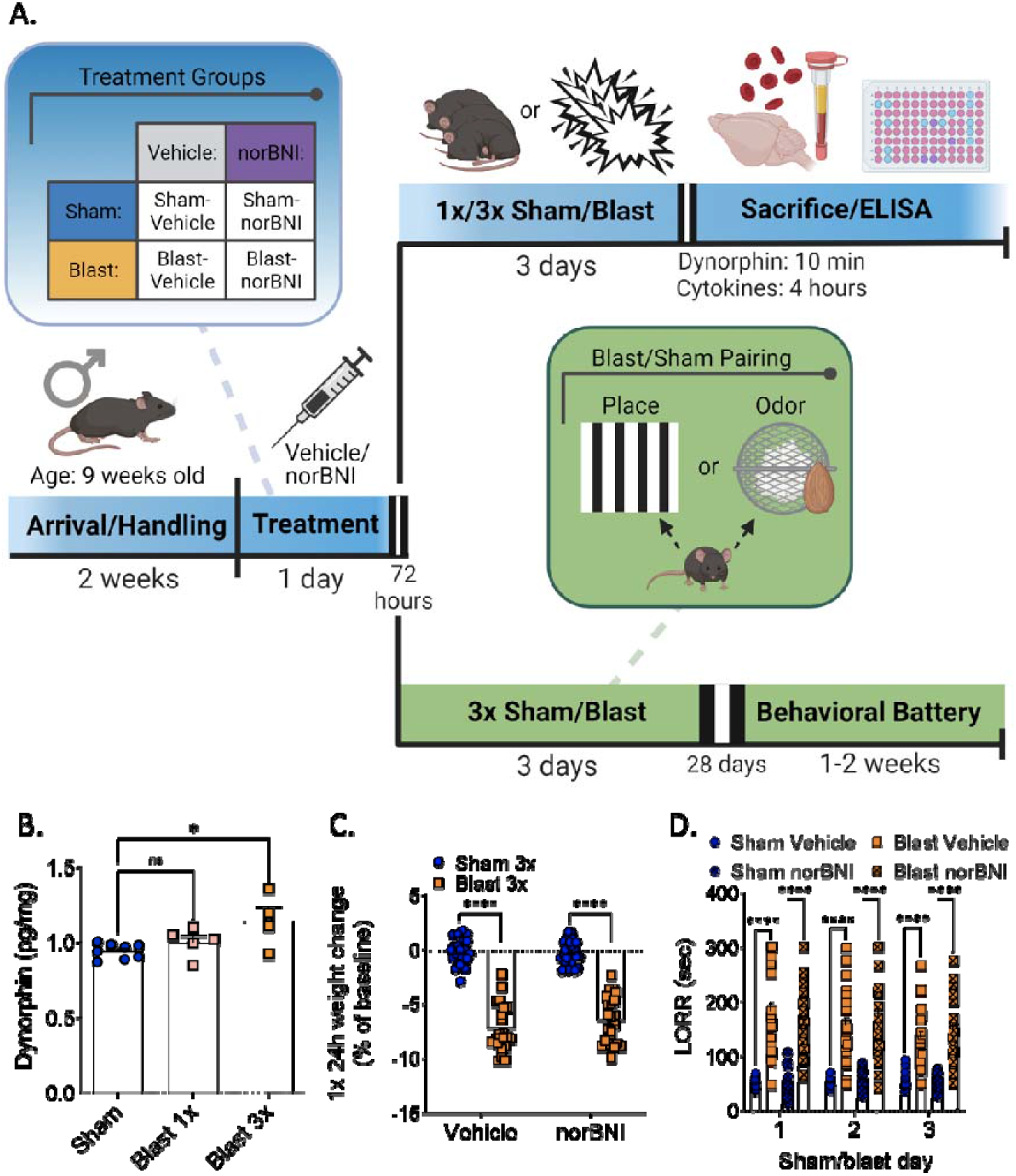
Timeline and acute effects of blast trauma. (A) Master timeline of experimental study. (B) Dynorphin A-like immunoreactivity is increased in the brain after 3x but not 1x blast. Oneway ANOVA. Tukey’s post-hoc analysis. (C) norBNI does not block acute weight loss. Two-way ANOVA. Bonferroni-Šídák post-hoc analysis. (D) norBNI does not block LORR delay. Two-way RM ANOVA. Tukey’s post-hoc analysis.

#### Dynorphin

Animals (n’s: sham: 8, blast 1x: 5, sham 3x: 4) were sacrificed 10 minutes following final sham/blast exposure brain collected for downstream biochemical analysis (Figure 1A). Brains were homogenized with ice cold lysis buffer as above. Dynorphin A-like immunoreactivity was assayed using a Dynorphin A enzyme immunoassay (EIA) kit (Raybiotech, Peachtree Corners, GA) as per manufacturer protocol (16-fold dilution) (Qi et al., 2013; Choi et al., 2021). The assay is a quantitative competition-based ELISA on a 96-well strip plate and Dynorphin Alike immunoreactivity is determined using a standard curve. Detection range: 0.1-1,000 ng/ml; sensitivity: 0.3 ng/ml. Measurements were completed by reading absorbance at 450 nm.

### Behavioral Battery

Behavioral testing was conducted one month post sham/blast exposure and consisted of two testing batteries each conducted in separate groups of mice over the course of one week (one test paradigm per day; Figure 1A). Each mouse only completed one of the testing batteries as follows: 1) marble burying, place conditioning, acoustic startle or 2) rotarod, odorant conditioning, photophobia reward conflict assay). The order of behavioral tests for each battery was performed from least to most stressful to decrease carryover distress from one behavior to the next. Prior to each day of testing, animals were placed in the testing room for at least 30 minutes to allow for acclimation. Cohort 1 n’s: sham vehicle: 17, sham norBNI: 19, blast vehicle: 17, blast norBNI: 15; Cohort 2 n’s: sham vehicle: 14, sham norBNI: 15, blast vehicle: 15, blast norBNI: 12.

### Cohort 1 (Figure 4A)

#### Marble burying

Animals were allowed to explore an open field in a clean rat cage, 259mm x 476mm x 209mm, 909 cm^2^ floor space) filled with 5 cm of bedding (kernels) and 18 marbles for 30 min. Marbles were counted as buried if at least two-thirds of the height of the marble was covered with bedding.

#### Place Conditioning Paradigm

A balanced-three compartment conditioning apparatus was used as described previously (Schindler et al., 2021a). One week prior to sham/blast exposure, animals were pre-tested by placing individual animals in the small central compartment and allowed exploration of the entire apparatus for 20 min. Mice were then counterbalanced to gray or vertical black and white stripes for their morning pairing based on initial side bias. Animals showing side bias were excluded. Sham/blast exposure occurred the following week over three consecutive days. On each day of exposure, in the morning, animals were placed in their AMpairing chamber containing distinct visual cues for 10 minutes and then were immediately given a sham exposure. In the afternoon of the same day, animals were placed in their PM-pairing chamber containing a different set of distinct visual cues for 10 min and then were immediately given a blast or sham exposure (depending on group assignment). Place conditioning was assessed at one month following exposure by allowing each mouse to roam freely in all three compartments. Time spent in each compartment was recorded and analyzed using Anymaze (Stoelting, Wood Dale, IL). Place conditioning scores were calculated by subtracting time spent in the PM paired compartment from time spent in the AM paired compartment.

#### Acoustic Startle

Conducted using SR-LAB acoustic startle boxes (San Diego Instruments, San Diego CA). Following a 5-min acclimation period, startle habituation testing consisted of 50 trials of 120-dB pulses delivered with an inter-trial interval of 7-23 s. Blast exposure can result in hearing loss in rodents (Mao et al., 2012), which also occurs in blast-exposed Veterans (Oleksiak et al., 2011). Therefore, we use the within subject analysis approaches of startle habituation in an attempt to mitigate potential confounds of hearing loss on startle outcome measures and interpretation. Startle habituation was expressed as the ratio of the average startle response in the last 10 trials to the average startle response in the first 10 trials (1.0 = no habituation).

### Cohort 2 (Figure 5A)

#### Rotarod

A rotarod apparatus (San Diego Instruments, San Diego, CA) was used to measure motor coordination and balance. The behavior was assessed over two days: acclimation and test day. On the acclimation day, three trials were used at the speed of 16 rpm. On the test day, five trials were used, each with a different set speed of rotation (in order of 8, 16, 24, 32, and 40 rpm). For each trial, the rod would accelerate to a set speed over 120s, remain at the speed for an additional 30 s, and finally decelerate back to 0 rpm over 30 s. The latency to fall off the rotarod was recorded for each trial and area under the curve (AUC) across all five trials was calculated.

#### Odorant Conditioning Paradigm

On each day of sham/blast exposure, animals received a stainless-steel mesh tea-ball (Amazon) containing one quarter Nestletts (PharmaServ, Framingham, MA) with 30 μl of imitation almond extract (Kroger, Cincinnati, OH) in their home cage five minutes prior to sham/blast exposure. The Nestletts and scents were refreshed on each subsequent sham/blast exposure and the tea-ball remained in place until 24 hours following final sham/blast exposure. One month following exposure, animals were tested for odorant-conditioning in a Plexiglas T-maze (66 cm long x 40 cm wide x 15 cm high) with a tea-ball containing one-quarter Nestlett with 30 μl imitation almond odorant cue placed in the left arm of the maze and a tea-ball containing one-quarter Nestlett with no odor placed in the opposite arm of the maze. Animals were placed in the long arm of the T-maze and given 5 min to explore the entire maze. Distance traveled and time spent in each of the two distal ends of the short arms were recorded and analyzed using Anymaze (Stoelting, Wood Dale, IL). Aversion is calculated as (scent/(scent + no scent).

#### Photophobia reward conflict assay

The photophobia assay was conducted in an Orofacial Pain Assessment Device (OPAD) using Anymaze acquisition and analysis software (Stoelting, Wood Dale, IL), and consisted of a choice between self-administration of a reward vs. exposure to a bright light. Mice were food-restricted one week prior and throughout the duration of testing. In order to decrease neophobia during training, mice were given ad libitum access to a sweetened condensed milk solution (1:2 solution of sweetened condensed milk in water (autoclaved, obtained from the animal facility) in their home cage for 24 hours. Animals were then trained to self-administer the milk solution from a sipper tube over 6 sessions (one session per day) of 9 minutes each. A lamp was positioned over the milk bottle but the light was not turned on during training sessions. The metal sipper tube with milk solution is attached to a stand and placed in a position where the spout is accessible to rodent, which enables the recording of individual licks as well as total lick duration. The general room lighting was maintained at 85 lux throughout. Testing occurred on the 7th session and consisted of an initial 3 minutes of no light followed by 3 minutes of a bright light (1000 lux) turned on directly above the sipper tube location, and then a final 3 minutes with no light. Photophobia ratio was calculated by dividing the duration of licking during the 3 minutes of light on by the duration of licking during the initial 3 minutes with no light.

### Data Analysis

As appropriate, data were analyzed using: (i) standard one-way analysis of variance (ANOVA), followed by Tukey’s Post-hoc tests; (ii) two-way (between/within subjects design) ANOVA, followed by Bonferroni-Šídák or Tukey’s Post-hoc tests (repeated measures (RM) where appropriate). Reported significant p values denote two-tailed probabilities of p ≤ 0.05 and non-significance (n.s.) indicates p > 0.05. Data were excluded in behavioral testing if the data point was equal to or greater than two standard deviations from the group mean. Statistical analyses were conducted using custom Python applications and Graph Pad Prism 4.0 (GraphPad Software, Inc., La Jolla, CA).

## RESULTS

Using well-established blast methods (Schindler et al., 2017; Schindler et al., 2021a; Schindler et al., 2021b; Baskin et al., 2021; Logsdon et al., 2018; Logsdon et al., 2020; Meabon et al., 2016), male C57Bl/6 adult mice were exposed to one (1x) or three (3x) blast BOPs using a pneumatic shock tube delivering a peak static pressure of 19.59 +/- 1.41 psi. Sham control mice were anesthetized equivalently to blast-exposed animals (see Materials and Methods). Figure 1a represents a general timeline of experiments performed in this study. Additional detail regarding cohorts undergoing each behavioral battery is shown in Figure 4A and 5A.

### Repetitive but not single blast exposure increases brain dynorphin A-like immunoreactivity, but KOR antagonism does not prevent blast-induced loss of righting reflex or acute weight loss

Corticosterone levels are increased in rodents after blast exposure (Schindler et al., 2021a; Zuckerman et al., 2018) and male mice demonstrate blast-induced aversion (Schindler et al., 2021a), highlighting a potential role for the dynorphin/KOR system. We first tested whether single or repetitive (3X) blast exposure (one exposure per day) resulted in changes to brain dynorphin A-like immunoreactivity levels using an immunoassay in whole brain lysate. We found that repetitive, but not single, blast exposure was necessary to elicit increased whole brain dynorphin A-like immunoreactivity levels 10 minutes post blast-exposure (one-way ANOVA: F[2,14]=5.146, p=0.021, Tukey’s; n=5-8) (Figure 1B).

Since dynorphin has been associated with stress exposure, we next investigated whether the long-acting receptor-inactivating KOR antagonist, norBNI, affects acute outcomes following sham or blast exposure. norBNI and saline vehicle was systemically administered to both blast and sham mice (see Figure 1a) three days prior to exposure. On exposure days, we measured weight change and LORR immediately after sham/blast as measures of acute physical stress response. We found that norBNI did not prevent weight loss 24-hours post-blast (two-way ANOVA: interaction effect F[1,86]=1.042, p=0.310, main effect of treatment F[1,86]=330.4, p<0.0001, main effect of drug F[1,86]=0.759, p=0.386, Bonferroni-Šídák; n=21-24) (Figure 1C) nor prevent the blast-induced LORR (two-way RM ANOVA: time x treatment effect F[6,172]=2.480, p=0.025, main effect of group F[3,86]=69.00, p<0.0001, main effect of time F[1.911,171.2]=1.976, p=0.142, Tukey’s; n=21-24) (Figure 1D).

### KOR antagonism blocks most cytokines acutely after repetitive blast exposure

Since cytokine release is commonly associated with blast trauma (Ruiecki et al., 2020; Gill et al., 2014) and dynorphin/KOR plays a role in inflammation and negative affect (Alicea et al., 1995; Bruchas et al., 2010; Zeng et al., 2020), we next tested whether KOR antagonism at the time of blast exposure blocks blast-induced cytokine changes in both brain and blood serum collected 4 hours post-exposure using an established 23-plex immunoassay (Figure 1A). The 23-plex assays for IL-1a, IL-1b, IL-2, IL-4, IL-5, IL-6, IL-7, IL-9, IL-10, IL-12(p40), IL-12(p70), IL-13, IL-15, IL-17, G-CSF, GM-CSF, INF-y, IP-10, MKC, MCP-1, MIP-1a, MIP-1b, MIP-2, RANTES, TNFa. A subset of the 23 analytes fell below the level of detection (detailed below) and were excluded from further analysis. These data are in agreement with the mild nature of our blast exposure paradigm as previously reported (Ruiecki et al., 2020; Gill et al., 2014). All statistics for this assay are detailed in Table 1 and Table 2 for serum and brain respectively. All analyses used two-way ANOVAs and post-hoc Bonferroni-Šídák.

**Table 1.** Serum Cytokine Statistics Statistical analysis of cytokine levels in blood serum. All analyses used two-way ANOVA and Bonferroni-Šídák post-hoc analysis showing p-values.

| Table 1: Serum Cytokine Statistics |  |  |  |  |  |  |
| --- | --- | --- | --- | --- | --- | --- |
| Analyte | Interaction F(df) | Interaction P | Treatment F(df) | Treatment P | Drug F(df) | Drug P |
| IL-1a | F[1,37] = 0.0622 | 0.8044 | F[1,37] = 0.4667 | 0.4987 | F[1,37] = 4.029 | 0.0521 |
| IL-5 | F[1,33] = 3.211 | 0.0823 | F[1,33] = 66.26 | <0.0001 | F[1,33] = 4.107 | 0.0509 |
| IL-6 | F[1,33] = 25.27 | <0.0001 | F[1,33] = 128.4 | <0.0001 | F[1,33] = 15.49 | 0.0004 |
| G-CSF | F[1,33] = 5.129 | 0.0302 | F[1,33] = 26.77 | <0.0001 | F[1,33] = 0.0033 | 0.9544 |
| IP-10 | F[1,36] = 1.409 | 0.243 | F[1,36] = 13.63 | 0.0007 | F[1,36] = 4.530 | 0.0402 |
| MKC | F[1,33] = 13.57 | 0.0008 | F[1,33] = 69.78 | <0.0001 | F[1,33] = 4.587 | 0.0397 |
| MCP-1 | F[1,35] = 0.3542 | 0.5556 | F[1,35] = 29.38 | <0.0001 | F[1,35] = 0.0068 | 0.9345 |
| Bonferroni-Šídák post-hoc analysis |  |  |  |  |  |  |
| Analyte | SV vs. BV. | SN vs. BN | SV vs. SN | BV vs. BN | SV vs. BN | BV. vs. SN |
| IL-1a | 0.9913 | 0.9997 | 0.7906 | 0.5126 | 0.9334 | 0.3124 |
| IL-5 | <0.0001 | <0.0001 | >0.9999 | 0.0492 | 0.0015 | <0.0001 |
| IL-6 | <0.0001 | 0.0001 | 0.9713 | <0.0001 | <0.0001 | <0.0001 |
| G-CSF | 0.0002 | 0.1603 | 0.5254 | 0.538 | 0.0057 | 0.0048 |
| IP-10 | 0.02 | 0.2843 | 0.1666 | 0.9831 | 0.002 | 0.8327 |
| MKC | <0.0001 | 0.0068 | 0.8776 | 0.0011 | 0.0011 | <0.0001 |
| MCP-1 | 0.0208 | 0.0002 | 0.9981 | 0.9994 | 0.0048 | 0.0018 |

**Table 2.** Brain Cytokine Statistics Statistical analysis of cytokine levels in brain lysate. All analysis used two-way ANOVA and Bonferroni-Šídák post-hoc analysis showing p-values.

| Table 2: Brain Cytokine Statistics |  |  |  |  |  |  |
| --- | --- | --- | --- | --- | --- | --- |
| Analyte | Interaction F(df) | Interaction P | Treatment F(df) | Treatment P | Drug F(df) | Drug P |
| IL-1a | F[1,21] = 8.038 | 0.0099 | F[1,21] = 3.223 | 0.087 | F[1,21] = 6.292 | 0.0204 |
| IL-6 | F[1,21] = 8.823 | 0.0073 | F[1,21] = 2.635 | 0.1194 | F[1,21] = 12.51 | 0.002 |
| IL-9 | F[1,21] = 6.426 | 0.0193 | F[1,21] = 5.718 | 0.0262 | F[1,21] = 10.72 | 0.0036 |
| IL-13 | F[1,21] = 13.96 | 0.0012 | F[1,21] = 6.627 | 0.0177 | F[1,21] = 2.730 | 0.1133 |
| IL-15 | F[1,21] = 8.788 | 0.0074 | F[1,21] = 0.4111 | 0.5284 | F[1,21] = 6.226 | 0.021 |
| IL-17 | F[1,21] = 6.958 | 0.0154 | F[1,21] = 4.310 | 0.0504 | F[1,21] = 1.663 | 0.2112 |
| G-CSF | F[1,21] = 13.49 | 0.0014 | F[1,21] = 23.16 | <0.0001 | F[1,21] = 3.591 | 0.0719 |
| INF-γ | F[1,21] = 27.88 | <0.0001 | F[1,21] = 7.497 | 0.0123 | F[1,21] = 4.560 | 0.0447 |
| IP-10 | F[1,21] = 5.875 | 0.0245 | F[1,21] = 27.45 | <0.0001 | F[1,21] = 3.743 | 0.0666 |
| MKC | F[1,21] = 0.7477 | 0.397 | F[1,21] = 0.1538 | 0.6989 | F[1,21] = 0.0077 | 0.9309 |
| MCP-1 | F[1,21] = 7.305 | 0.0132 | F[1,21] = 12.17 | 0.0022 | F[1,21] = 5.254 | 0.0323 |
| RANTES | F[1,21] = 12.65 | 0.0019 | F[1,21] = 15.61 | 0.0007 | F[1,21] = 3.828 | 0.0638 |
| Bonferroni-Šídák post-hoc analysis |  |  |  |  |  |  |
| Analyte | SV vs. BV. | SN vs. BN | SV vs. SN | BV vs. BN | SV vs. BN | BV. vs. SN |
| IL-1a | 0.0397 | >0.9999 | >0.9999 | 0.0094 | >0.9999 | 0.0273 |
| IL-6 | 0.0419 | >0.9999 | >0.9999 | 0.0014 | >0.9999 | 0.0062 |
| IL-9 | 0.0254 | >0.9999 | >0.9999 | 0.0044 | >0.9999 | 0.0025 |
| IL-13 | 0.003 | >0.9999 | 0.834 | 0.0087 | >0.9999 | 0.0311 |
| IL-15 | 0.173 | 0.5034 | >0.9999 | 0.0078 | >0.9999 | 0.185 |
| IL-17 | 0.035 | >0.9999 | >0.9999 | >0.0860 | >0.9999 | 0.1291 |
| G-CSF | 0.0001 | >0.9999 | >0.9999 | 0.0065 | 0.3633 | 0.0004 |
| INF-γ | 0.0002 | 0.3617 | 0.1826 | 0.0003 | >0.9999 | 0.0102 |
| IP-10 | 0.0004 | 0.2353 | >0.9999 | 0.0446 | 0.2133 | 0.0002 |
| MKC | >0.9999 | >0.9999 | >0.9999 | >0.9999 | >0.9999 | >0.9999 |
| MCP-1 | 0.0036 | >0.9999 | >0.9999 | 0.0164 | >0.9999 | 0.0021 |
| RANTES | 0.0005 | >0.9999 | >0.9999 | 0.0071 | >0.9999 | 0.0017 |

In serum, only seven cytokines were detected at appreciable levels: IL-1α, IL-5, IL-6, G-CSF, IP-10, MKC (CXCL1), and MCP-1. norBNI pretreatment had differential effects on specific cytokines in the serum. Neither repetitive blast exposure nor KOR antagonist treatment altered IL-1α levels (Figure 2A). Repetitive blast exposure significantly increased IL-5, IL-6, G-CSF, IP-10 and MKC in blast when compared to the sham controls, and norBNI attenuated those effects (Figure 2B-2F). norBNI did not affect blast-induced MCP-1 increase (Figure 2G).

**Figure 2.**
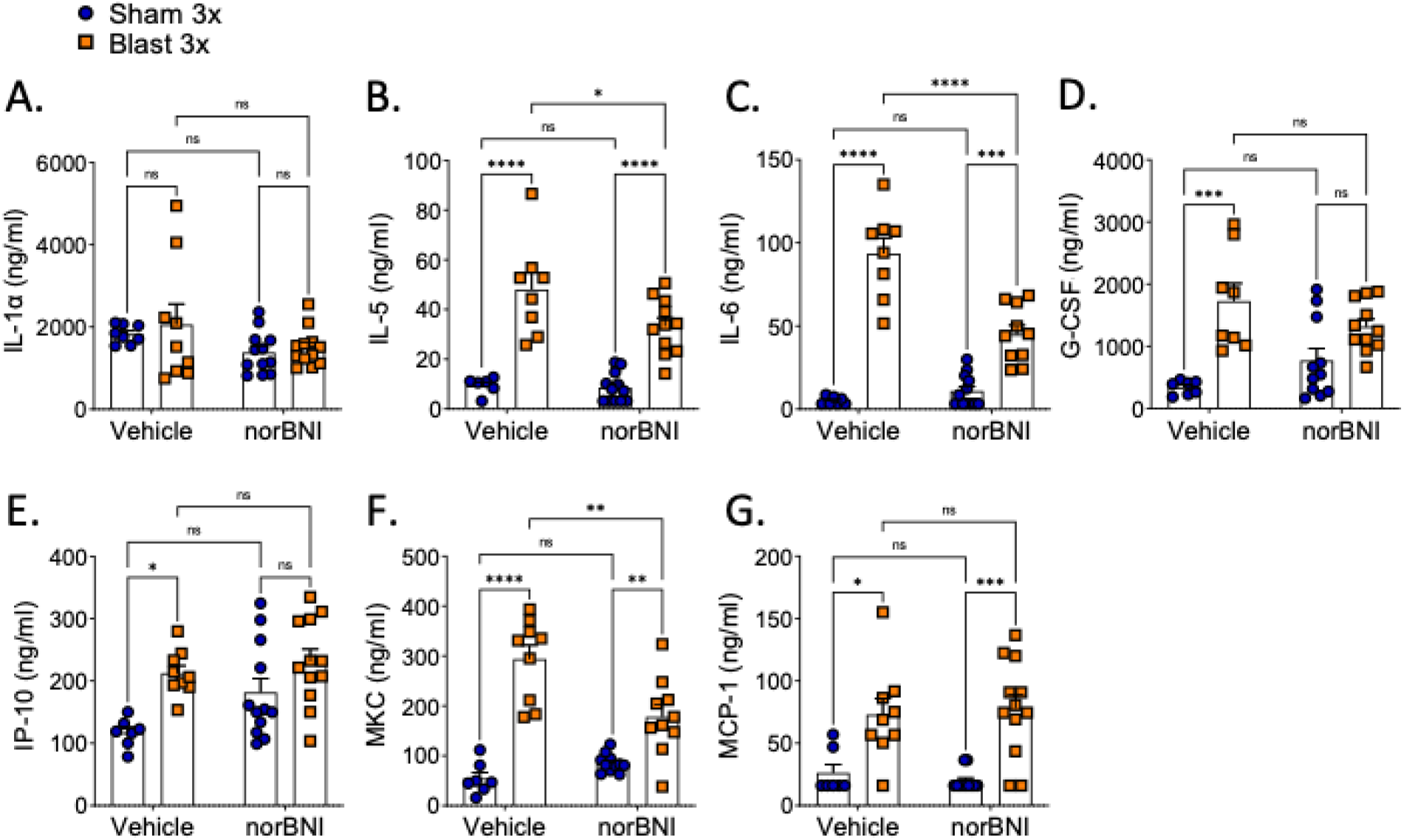
norBNI pretreatment blocked some but not all inflammatory cytokines in blood serum. norBNI nor blast exposure alter (A) IL-1α levels. norBNI attenuated blast-induced (B) IL-5, (C) IL-6, and (F) levels. norBNI did not block blast-induced (D) G-CSF and (E) IP-10 levels and there is no difference between norBNI groups. (G) norBNI did not block blast-induced MCP-1 levels. (A-G) Two-way ANOVA. Bonferroni-Šídák post-hoc analysis. *p ≤ 0.05, **p ≤ 0.01, ***p ≤ 0.001, ****p ≤ 0.0001, ns = not significant. Values represent mean ± SEM.

In brain lysates, 12 cytokines were detected: IL-1α, IL-6, IL-9, IL-15, IL-17, G-CSF, INF-γ, IP-10, MKC, MCP-1, and RANTES (Figure 3A-L). Compared to serum, blast and norBNI effects were more robust and consistent in the brain. All cytokines detected except IL-15 (Figure 3E) and MKC (Figure 3J) had significantly increased levels in the blast vehicle group as compared to the sham vehicle group (trend for increase in IL-15 following blast). When compared to the vehicle blast group, norBNI blocked all blast-induced changes except for MKC (Figure 3J). Of the cytokines analyzed, IL-6 was the only cytokine whose increase following repetitive blast exposure was blocked by norBNI in both serum and brain.

**Figure 3.**
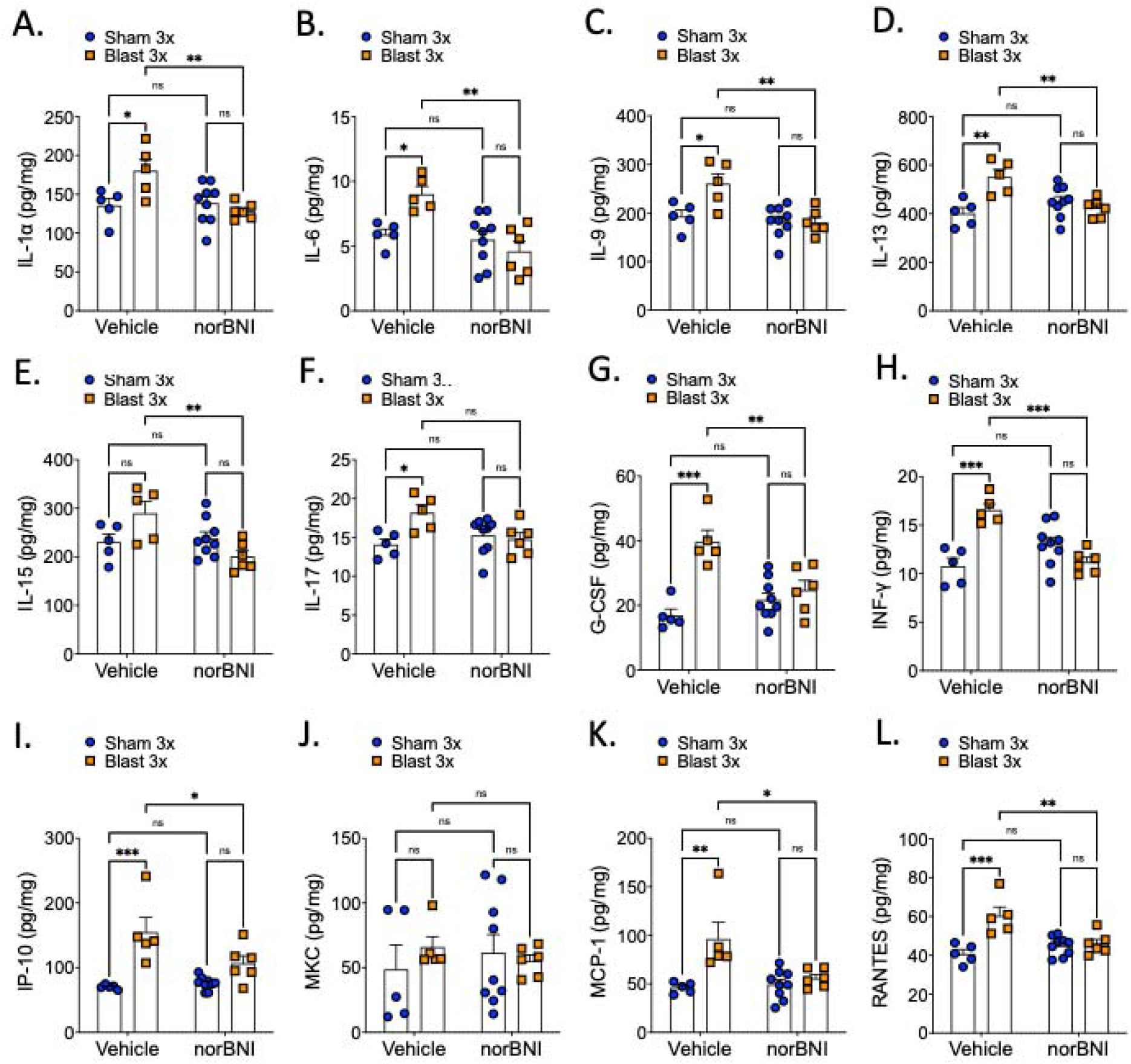
norBNI pretreatment blocked most but not all inflammatory cytokines in the brain. norBNI blocks blast-induced released in (A) IL-1a, (B) IL-6, (C) IL-9, (D) IL-13, (E) IP-15, (G) G-CSF, (H) INF-y, (I) IP-10, (K) MCP-1, and (L) RANTES, but not in (F) IL-17. (J) MKC levels are not affected by blast nor norBNI. (E) IP-15 does not increase in the vehicle blast group. (A-L) Two-way ANOVA. Bonferroni-Šídák post-hoc analysis. *p ≤ 0.05, **p ≤ 0.01, ***p ≤ 0.001, ****p ≤ 0.0001, ns = not significant. Values represent mean ± SEM.

### KOR antagonism blocks repetitive blast-induced anxiety-/aversive-like behavior and photosensitivity but not ambulatory deficit

Separate sets of mice were tested in two sets of behavioral measures. In the first group of mice, a discrete environmental stimulus was presented for 10 minutes prior to sham/blast exposure. Behavioral testing then occurred one month following repetitive sham or blast exposure and included marble burying (anxiety/compulsivity), place conditioning (aversion), and acoustic startle (habituation) (Figure 4A). norBNI pretreatment blocked aversion to paired visual cues (two-way ANOVA: interaction effect F[1,41]=2.499, p=0.122, main effect of treatment F[1,41]=6.943, p=0.012, main effect of drug F[1,41]=3.667, p=0.063, Bonferroni-Šídák; n=10-12) (Figure 4B), and did not have an effect on distance traveled (two-way ANOVA: interaction effect F[1,41]=0.055, p=0.815, main effect of treatment F[1,41]=0.577, p=0.452, main effect of drug F[1,41]=1.251, p=0.270, Bonferroni-Šídák; n=10-12) (Figure 4C). Therefore, differences in time spent in the blast-paired environment were not attributable to locomotor differences. Likewise, norBNI blocked the blast-induced increased in number of marbles buried (two-way ANOVA: interaction effect F[1,64]=4.580, p=0.036, main effect of treatment F[1,64]=2.370, p=0.129, main effect of drug F[1,64]=1.007, p=0.319, Bonferroni-Šídák; n=15-19) (Figure 4D) and inhibited acoustic startle habituation (two-way ANOVA: interaction effect F[1,51]=6.194, p=0.016, main effect of treatment F[1,51]=2.643, p=0.11, main effect of drug F[1,51]=2.161, p=0.148, Bonferroni-Šídák; n=12-15) (Figure 4E).

**Figure 4.**
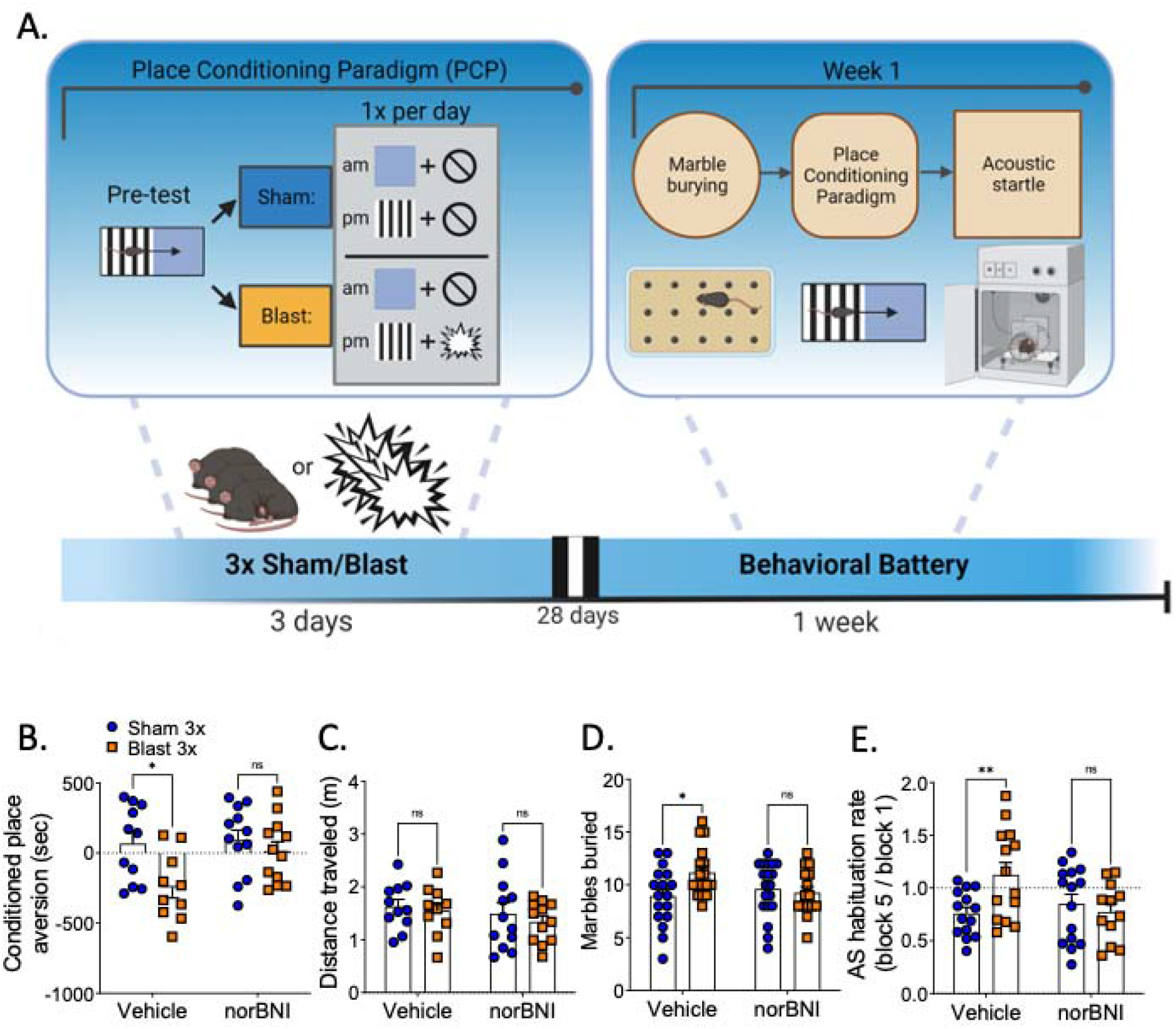
norBNI blocks blast effect in anxiety- and aversive-like behavior in PCP. (A) Timeline of behavioral battery of cohorts undergoing PCP. (B) norBNI blocks aversive-like behavior in PCP. (C) norBNI does not block locomotion in PCP. (D) norBNI blocks blast effect in number of marbles buried. (E) norBNI blocks delayed habituation rate blast effect in acoustic startle response. (B-E) Two-way ANOVA. Bonferroni-Šídák post-hoc analysis. *p ≤ 0.05, **p ≤ 0.01, ns = not significant. Values represent mean ± SEM.

For the second group of mice, sham/blast exposure was paired with a liquid almond extract liquid absorbed onto a Nestlet. The Nestlet with almond extract was enclosed in a tea ball strainer and placed in the cage for 5 minutes before sham or blast exposures (Figure 5A). One month following repetitive sham or blast exposure, mice were tested in order listed: OCP (aversion), rotarod (motor coordination), and the photophobia reward conflict (photosensitivity) as sensitivity to light has been reported in blast TBI patients (Truong et al., 2014, Callahan & Lin 2018). In line with results from the place conditioning experiments, norBNI blocked aversion to blast-paired odor (two-way ANOVA: interaction effect F[1,50]=4.160, p=0..047, main effect of treatment F[1,50]=4.214, p=0.043, main effect of drug F[1,50]=0.111, p=0.740, Bonferroni-Šídák; n=12-16) (Figure 5B) but did not affect distance traveled (two-way ANOVA: interaction effect F[1,50]=0.473, p=0.495, main effect of treatment F[1,50]=0.328, p=0.569, main effect of drug F[1,50]=0.291, p=0.592, Bonferroni-Šídák; n=12-16) (Figure 5C). Conversely, norBNI did not block or decrease blast-induced deficits on the accelerating rotarod (two-way ANOVA: interaction effect F[1,74]=2.419, p=0.124, main effect of treatment F[1,74]=23.91, p<0.0001, main effect of drug F[1,74]=1.031, p=0.313, Bonferroni-Šídák; n=19-20) (Figure 5D). Following these behaviors, mice had their food access restricted and were given access to a sweetened condensed milk solution in a modified OPAD for 9 minutes per day (See Methods). On the final test day, a bright light was lit directly over the access point to the milk solution at the 6-minute mark. Remarkably, we found norBNI blocked increased blast-induced aversion to light (two-way ANOVA: interaction effect F[1,50]=6.733, p=0.012, main effect of treatment F[1,50]=0.562, p=0.457, main effect of drug F[1,50]=0.767, p=0.385, Bonferroni-Šídák; n=12-15) (Figure 5E).

**Figure 5.**
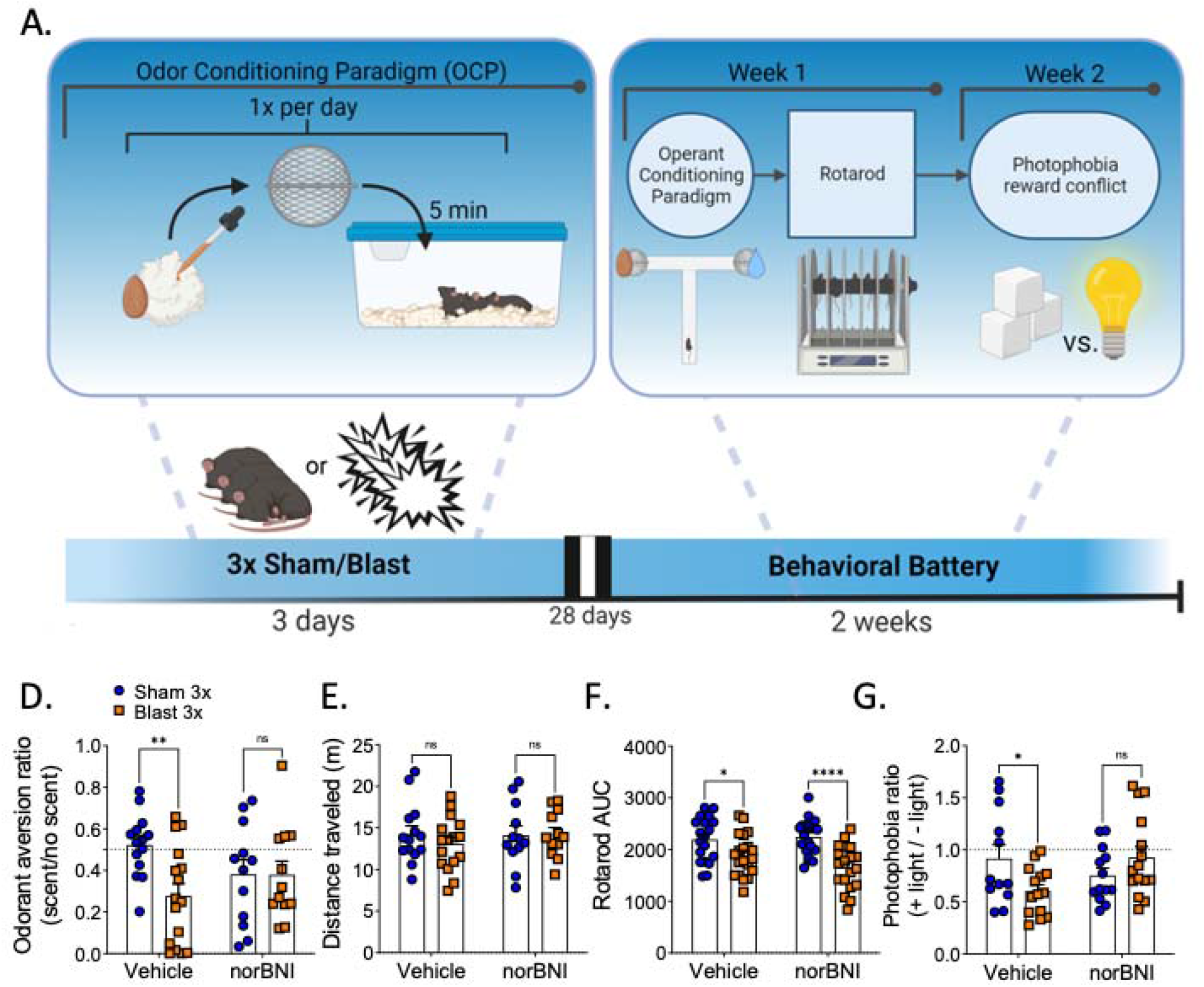
norBNI blocks blast effect in light sensitivity- and aversive-like behavior in OCP. (A) Timeline of behavioral battery of cohorts undergoing OCP. (B) norBNI blocks aversive-like behavior in OCP. (C) norBNI does not block locomotion in OCP. (D) norBNI does not block blast effect in latency to fall in rotarod. (E) norBNI blocks blast effect in time spent away from light. (BE) Two-way ANOVA. Bonferroni-Šídák post-hoc analysis. *p ≤ 0.05, **p ≤ 0.01, ns = not significant. Values represent mean ± SEM.

## DISCUSSION

In this study, we uncovered a critical role for repetitive blast-induced dynorphin/KOR activation in mediating discrete aspects of (neuro)-inflammation and behavioral dysfunction at acute and chronic time points in male mice. The majority of all TBIs in the US are mild (CDC 2003; WHO 2006; DOD 2021) and can result in significant adverse outcomes and decreased quality of life. However, currently there are no approved prophylactic or therapeutic treatment approaches for adverse outcomes following blast TBI. While the dynorphin/KOR system has been implicated in severe impact TBI (McIntosh et al., 1987; Redell et al., 2002; Hussain et al., 2012), there has been no investigation of this receptor system in models of impact or blast mTBI. Here, we present novel evidence for an increase in brain-wide dynorphin A-like immunoreactivity acutely following repeated, but not single blast exposure. These results are consistent with previous findings that demonstrate only repetitive blast exposure resulted in aversion to blast-paired cues (Schindler et al., 2021a). Furthermore, here we extend on our previous finding by showing that systemic administration of the selective long-lasting KOR antagonist norBNI blocked or decreased blast-induced cytokine changes in serum and brain samples and discrete aspects of chronic behavioral dysfunction. Together these results highlight the dynorphin/KOR system as a mediator of blast-induced dysfunction and support further research aimed at establishing this system as a potential prophylactic and/or therapeutic target for adverse outcomes following repetitive blast trauma.

In this study, we focused our efforts on norBNI, a selective long-acting KOR-antagonist lasting up to 21 days in rodents (Bruchas & Chavkin, 2010; Bruchas et al., 2010; Bruchas et al., 2007; Chavkin et al., 2019; Schattauer et al., 2017). While administration of a KOR antagonist had effects on acute cytokine levels and chronic behavioral dysfunction, KOR antagonism did not block LORR immediately following blast exposure nor acute weight loss, suggesting that these adverse outcomes are mediated by other systems and processes. Likewise, KOR antagonism blocked or altered acute cytokine changes following repetitive blast exposure in the brain more robustly than in blood serum. Together, these results demonstrate the ability of this pharmacological approach to isolate and link discrete pathophysiological and behavioral outcomes associated with mTBI, which can in turn inform downstream translation and precision medicine efforts.

Pretreatment with norBNI blocked blast-induced IL-6 increase, which may offer neuroprotection in acute injury but has also been correlated with health risks chronically in both TBI and PTSD (Monsour et al., 2022). The differential effect between the brain and serum cytokine levels may be due to alterations of blood-brain-barrier (BBB) permeability or differing dynorphin/KOR mechanisms in the periphery versus the brain (Bruchas et al., 2010; Cahill et al., 2014; Best et al., 2022). While there is a lack of research investigating dynorphin/KOR involvement in BBB permeability, KOR may be implicated in BBB integrity through its actions on the HPA axis and/or inflammatory response. Despite having acutely increased cortisol (Cernak et al., 1999), SMs have below baseline levels of cortisol at more chronic timepoints (Yehuda et al., 2009) and TBI (Agha et al., 2007). Additionally, glucocorticoids may play an important role for BBB recovery in blast trauma by tightening the barrier resulting in decreased peripheral immune invasion and inflammation (Hue et al., 2015). Furthermore, blast-induced activation of the dynorphin/KOR system may influence (neuro)-inflammation and BBB dysfunction through priming stress-stimulated immune cells (Sapolsky et al., 1986; Tapp et al., 2019). Importantly, systemic administration of norBNI can pass the BBB (Munro et al., 2012; Patkar et al., 2013), therefore, KOR antagonism may provide protective benefits via mechanisms outside of BBB disruption and/or systemic inflammation.

Moving forward, a detailed examination of dynorphin and cytokine changes post blast in discrete brain regions (e.g., amygdala, cortex, striatum, cerebellum) is warranted as results here detail whole brain levels, which may mask more discrete changes within specific brain regions. These efforts are ongoing and in parallel with immunohistochemical efforts to understand a potential role for dynorphin/KOR in blast-induced microglia and astrocyte dysfunction. Our results demonstrating an effect of systemic KOR antagonism on adverse blast outcomes suggest that repetitive blast trauma results in release of dynorphin and activation of KORs. Conversely, it is important to note that the current study did not directly assess dynorphin release, so while unlikely, the increase in dynorphin A-like immunoreactivity seen here could alternatively represent increased production and storage of dynorphin in intracellular vesicles. Experiments using subcellular fractionation techniques to isolate dense core vesicles (vesicles which release dynorphin peptide from neurons) and/or in vivo imaging of a genetically encoded dynorphin sensor (Abraham et al., 2021) could also increase understanding of how and where blast exposure impacts dynorphin release and cytokine expression. Likewise, mass-spectrometric examination of prodynorphin gene products in extracellular fluid pre/post-blast might also lend important information regarding how blast trauma affects prodynorphin processing and release.

Depression and anxiety are common behavioral symptoms associated with blast trauma in Veterans and SMs, and there is ongoing debate regarding specific contributions of blast-induced mTBI vs. PTSD to these long-term outcomes (Hendrickson et al., 2018). Here, we provide evidence that the dynorphin/KOR system encodes aversive- and anxiety-like components of blast trauma. We also find that KOR antagonism blocked aversive-like behavior in a photophobia assay, suggesting this adverse outcome may be stress-mediated and associated with negative pain affect (Cahill et al., 2014). Conversely, here we did not find an effect of the dynorphin/KOR system on motor coordination skills. This is consistent with the proposed role of the dynorphin/KOR system in primarily mediating affective components of stress and trauma exposure (Van’t Veer & Carlezon 2013; Hang et al., 2015). As rotarod requires skill in balance, grip strength, and motor coordination, mechanisms outside of the dynorphin/KOR system may be mediating these effects (Mouzon et al., 2012). We have previously demonstrated a blast-induced decrease in rotarod performance that is mediated by nitric oxide synthase-dependent BBB permeability (Logsdon et al., 2018; Logsdon et al., 2020), and a relationship between dynorphin and nitric oxide has been previously suggested (Sharma & Alm 2002, Hu et al., 1999), but a role for dynorphin/KOR in BBB permeability nor NOS function post repetitive blast has not been examined.

Our results underscore the need for further investigation of the dynorphin/KOR system’s role in blast-induced trauma, including executive dysfunction and chronic pain. The dynorphin/KOR system interfaces with mesolimbic circuitry and can regulate dopamine dynamics, and we have previously reported aberrant phasic dopamine release in the nucleus accumbens and executive dysfunction in both mice and Veterans with blast mTBI (Schindler et al., 2017; Baskin et al., 2021; Escobar et al., 2020). Therefore, future directions should include investigation of dynorphin/KOR’s involvement in blast-induced dopamine dysfunction and adverse behavioral outcomes related to executive function. Additionally, the dynorphin/KOR system has been implicated in drug reward and substance use disorder, which is increased in SMs and Veterans following blast trauma and in mouse models of blast (Schindler et al., 2021b; Petriak et al., 2012; Miller et al., 2012). Lastly, we recognize the importance of assessing the potential contribution of sex as a biological variable in mechanisms underlying blast trauma outcomes (McCabe & Tucker 2020) as sex differences in dynorphin/KOR function have been previously reported (Chartoff & Mavrikaki 2015) and combat military occupational specialties are now available not only to males. In order to further increase translational relevance, additional blast modeling efforts are also warranted as Servicemembers’ blast exposure experience can vary widely in number of blast exposures (1-100s have been previously reported) and/or intervals between blasts (e.g., multiple blasts over a day or multiple days vs. multiple blasts from multiples deployments) (Huber et al., 2016; Peskind et al., 2011; Petrie et al., 2014).

## CONCLUSIONS

In conclusion, this study demonstrates a critical role for the dynorphin/KOR system in mediating discrete aspects of (neuro)-inflammation and behavioral dysfunction at acute and chronic time points following repetitive blast trauma in male mice. These results highlight the dynorphin/KOR system as a promising prophylactic and therapeutic target for overlapping comorbidities resulting from repetitive blast trauma and demonstrate KOR antagonists as a tool for increased mechanistic understanding of blast-induced psychopathology. While an estimated 400,000 Veterans have experienced blast mTBI (Hoge et al., 2008; Tanielian & Jaycox 2008; DOD 2021), to date there are currently no approved pharmacological agents for prevention/treatment of adverse outcomes. KOR antagonists are currently under clinical trial investigation in the civilian population for treatment of non-blast stress-related disorders (Carlezon & Krystal 2016), but the potential of these drugs in a blast setting has not been investigated. Therefore, increased research devoted to prevention/treatment efforts with KOR antagonists in animal models and in clinical settings is warranted and has a high likelihood of success.

## ABBREVIATIONS

ANOVA: analysis of variance
BBB: blood brain barrier
BOP: blast overpressure
HPA: hypothalamic-pituitary-adrenal
KOR: kappa opioid receptor
LORR: loss of righting reflex
mTBI: mild traumatic brain injury
norBNI: nor-binaltorphimine dihydrochloride
OCP: odorant conditioning paradigm
OIF/OEF/OND: Operation Iraqi Freedom/Operation Enduring Freedom/Operation New Dawn
OPAD: Orofacial Pain Assessment Device
PCP: place conditioning paradigm
PTSD: posttraumatic stress disorder
SM: servicemember

## DISCLAIMER

The views expressed in this scientific presentation are those of the author(s) and do not reflect the official policy or position of the U.S. government or Department of Veteran Affairs.

## DECLARATIONS

### Ethics approval and consent to participate

All animal experiments were conducted in accordance with Association for Assessment and Accreditation of Laboratory Animal Care guidelines and were approved by the VA Puget Sound Institutional Animal Care and Use Committee.

### Consent for publication

Not applicable.

### Availability of data and materials

The data in this study are available from the corresponding author upon reasonable request.

### Authors’ contributions

The work presented here was carried out in collaboration among all authors. AS, SJL, AL, and DG contributed to conception and design of the study. SJL, AL, MY, BB, and AS collected and analyzed data. SJL and AS wrote the first draft of the manuscript. All authors contributed to manuscript revision, read, and approved the final manuscript.

## Acknowledgements

We would like to thank Scott Ng Evans, Traci J. Weber, Cindy Pekow, DVM, Kari Koszdin, DVM, Monica, and Lena Strait-Bodeyfor considerable technical assistance and veterinary care.

